# PII1: a protein involved in starch initiation that determines granule number and size in Arabidopsis chloroplast

**DOI:** 10.1101/310003

**Authors:** Camille Vandromme, Corentin Spriet, David Dauvillée, Adeline Courseaux, Jean-Luc Putaux, Adeline Wychowski, Maud Facon, Christophe D’Hulst, Fabrice Wattebled

## Abstract

The initiation of starch granule formation is still poorly understood. However, soluble starch synthase 4 (SS4) appears to be a major component of this process since it is required to synthetize the correct number of starch granules in the chloroplasts of *Arabidopsis thaliana* plants. A yeast-2-hybrid screen allowed the identification of several putative SS4 interacting partners. We identified the product of *At4g32190* locus as a chloroplast-targeted PROTEIN INVOLVED IN STARCH INITIATION (named PII1). Arabidopsis mutants devoid of PII1 display an alteration of starch initiation process and accumulate, on average, one starch granule per plastid instead of the 5 to 7 granules found in plastids of wild-type plants. These granules are larger than in wild type and they remain flat and lenticular. *pii1* mutants display wild-type growth rates and accumulate standard starch amounts. Moreover, starch characteristics, such as amylopectin chain length distribution, remain unchanged. Our results reveal the involvement of PII1 in starch priming process in Arabidopsis leaves through interaction with SS4.

## Introduction

Starch is the main storage polysaccharide produced by plants. It accumulates as water-insoluble semi-crystalline granules in the chloroplast of photosynthetic organ cells or in the amyloplasts of storage organ cells (potato tubers, endosperm of cereal seeds). Starch is a mix of two structurally distinct α-glucan polymers, amylose and amylopectin in which glucose residues are linked in α(1→4) and branched in α(1→6). Amylopectin, the major fraction of starch, is moderately branched containing up to 6% of α(1→6) linkages while the frequency of branching of amylose is much below 1%.

Starch synthesis is a complex process that implies tens of proteins, enzymatically active or not, and each step is catalyzed by several genetically independent isoforms (D’Hulst *et al*., 2015). For instance, up to five starch-synthases catalyze the elongation of the α-glucan polymers by transferring the glucose residue from ADP-glucose to the non-reducing end of the molecules (Abel *et al*., 1996; Edwards *et al*., 1999; Delvallé *et al*., 2005; Zhang *et al*., 2005; Zhang *et al*., 2008; Crofts *et al*., 2017). The formation of the branch points and the control of their distribution in amylopectin is monitored by up to three branching enzymes that create α(1→6) bonds (Schwall *et al*., 2000; Blauth *et al*., 2001; Tanaka *et al*., 2004; Yao *et al*., 2004; Dumez *et al*., 2006; Nakamura *et al*., 2010; Regina *et al*., 2010; Tetlow, 2012) and by debranching enzymes (isoamylases and pullulanase) that hydrolyze some of them (Mouille *et al*., 1996; Myers *et al*., 2000; Delatte *et al*., 2005; Wattebled *et al*., 2005; Streb *et al*., 2008; Wattebled *et al*., 2008; Ferreira *et al*., 2017). This process induces the formation of a cluster-like structure of amylopectin responsible for its specific physicochemical properties (Pfister & Zeeman, 2016).

A major current challenge is to understand how the activity of these enzymes is controlled to generate new starch granules. It is now well established that enzymes such as branching enzymes and starch synthases are engaged in hetero-multimeric complexes (Tetlow *et al*., 2004; Hennen-Bierwagen *et al*., 2008; Tetlow *et al*., 2008; Hennen-Bierwagen *et al*., 2009; Ahmed *et al*., 2015; Crofts *et al*., 2015). However, the regulation of the formation of these complexes remains to be elucidated even if it is strongly suspected that the protein phosphorylation state is a key factor controlling protein-protein interaction (Liu *et al*., 2009; Liu *et al*., 2012; Makhmoudova *et al*., 2014; Subasinghe *et al*., 2014). Moreover, an increasing number of non-catalytic proteins have been described to be involved in starch metabolism with functions that are essential for correct starch synthesis or degradation (Seung *et al*., 2015; Feike *et al*., 2016; Seung *et al*., 2017).

One step of starch synthesis that remains poorly understood is the initiation of granule formation. This process is of prime importance since it defines, *in fine*, the number, the size, and the morphology of the starch granules. Arabidopsis accumulates on average 5 to 7 starch granules per plastid in mature leaves. This number is rather constant, implying a finely tuned regulation *in planta*, and depends on the chloroplast volume (Crumpton-Taylor *et al*., 2012; Crumpton-Taylor *et al*., 2013). It has been shown that starch synthase 4 (SS4) is a major factor affecting the priming of starch synthesis. Arabidopsis *ss4* mutants accumulate one (sometimes none, rarely two) starch granule(s) per chloroplast (Roldán *et al*., 2007). Interestingly, this reduction in the number of starch granules per plastid is accompanied by a modification of their shape. Wild-type (WT) granules are generally flat and lenticular with a diameter of 1-2 μm. Starch granules in the *ss4* mutant are larger (3-5 μm) and spheroidal (Roldán *et al*., 2007). The synthesis of the unique granule in the *ss4* mutant depends on the presence of another starch synthase: SS3. Indeed, starch synthesis collapses in the *ss3 ss4* double mutant (Szydlowski *et al*., 2009) and the synthesis of one starch granule observed in few chloroplasts are probably due to stochastic initiation events (Crumpton-Taylor *et al*., 2013).

SS4 is a protein composed of two distinct domains. The C-terminal moiety of the protein corresponds to the glycosyl-tranferase 5 (GT5) domain of the CAZy classification that is shared by all starch synthases (Coutinho *et al*., 2003). The N-terminal half of the protein is specific to SS4 and is essentially composed of coiled-coil motifs (Leterrier *et al*., 2008; Gámez-Arjona *et al*., 2014). These two domains have specific functions in granule formation. While the C-terminal part of the protein determines the number of initiation events, the N-terminal moiety is involved in protein localization and controls granule shape (Lu *et al*., 2018). Indeed, SS4 is not evenly distributed within the chloroplast but is associated with plastoglobules where it has been described to interact with fibrillins 1 (Gámez-Arjona *et al*., 2014; Raynaud *et al*., 2016). It was also reported that SS4 interacts with itself and with PTST2, a non-catalytic protein that, together with PTST3, was proposed to deliver a substrate allowing SS4 to initiate starch granule formation (Seung *et al*., 2017).

In this article, we report the identification of a new protein that physically interacts with SS4 and is involved in starch priming. This protein was named PII1 for “Protein Involved in starch Initiation” (*At4g32190*). Mutants lacking PII1 have a reduced number of larger starch granules compared to the wild type. The phenotype observed is not a strict phenocopy of that of the *ss4* mutant, because plant growth and starch granule morphology are unaltered in *pii1* mutant compared to wild type. We propose that PII1 is required for starch granule initiation by controlling the catalytic activity of SS4.

## Material and methods

### Plant material and growth conditions

The ULTImate Y2H™ was carried out by Hybrigenics-services (Paris, France) using SS4 as bait (amino acids 43 to 1040) against a library prepared from one-week-old seedlings. Among 125 millions interaction tested, 369, corresponding to 80 different proteins, were positives. Using the ChloroP algorithm prediction (Emanuelsson *et al*., 1999), we were able to select proteins with predicted chloroplast targeting peptides. We ended-up with six candidates among which PII1 (*At4g32190*) was selected (Table S1). Hybrigenics-services provides interaction results associated to a predicted biological score (PBS). This score indicates the interaction reliability and is divided in 6 different classes (A to F): A: very high confidence in the interaction. B: high confidence in the interaction. C: good confidence of interaction. D: moderate confidence of interaction. E: interaction involving highly connected prey domains. This class is subjected to non-specific interactions. F: experimentally proven artifacts.

*Arabidopsis thaliana* lines were obtained from NASC (Nottingham Arabidopsis Stock Centre; http://Arabidopsis.info; (Alonso *et al*., 2003)) or from the collection generated at URGV (INRA of Versailles; (Samson *et al*., 2002)). Wassilewskija (Ws) and Columbia (Col-0) lines were used as wild type references. T-DNA insertion lines used are: *pii1-1* (SALK_122445); *pii1-2* (FLAG_137A02); *ss4-1* (GABI_290D11); *ss4-2* (FLAG_559H08). Both *pii1-1* and *ss4-1* are in Columbia genetic background while *pii1-2* and *ss4-2* were generated in Ws genetic background. *ss4* alleles were already described in (Roldán *et al*., 2007). Oligonucleotides used for selection and RT-PCR experiments are described in Table S2.

Depending on experiments, plants were grown either in a greenhouse (16 h : 8 h, light : dark photoperiod at 23 °C during the day and 20 °C during the night, 150 μmol photon m^−2^ s^−1^) or in a growth chamber (16 h : 8 h, light : dark photoperiod at 23 °C during the day and 20 °C during the night, 120 μmol photon m^−2^ s^−1^). Seeds are incubated at 4 °C in 0.1 *%* Phytagel solution (w/v) during 3 days before being sown on peat-based compost.

The selection of homozygous mutant lines was performed by PCR amplification on genomic DNA according to standard procedures described in Wattebled *et al*, 2008. RNA were extracted from leaves harvested at the middle of the light phase using Nucleospin RNA plant kit from Macherey-Nagel following manufacturer instructions. 500 ng of RNA were used to complete RT-PCR amplification using the One-Step RT-PCR kit from Qiagen. To ensure that RNA extraction was correctly performed, we have systematically amplified, as a negative control, template without the step of retro transcription. Detailed primer sequences are listed in table S2.

The analysis of starch accumulation in roots was performed on plants grown under hydroponic conditions. The seeds were sterilized in ethanol 75% and stratified at 4 °C during 3 days. Seeds were then deposited on the Seedholder (araponics^®^) completed with Murashige and Skoog medium and 0.8% plant agar (Duchefa Biochemie). Roots develop in a culture solution (1.1 mM MgSO_4_, 2mM KNO_3_, 805 μM Ca(NO_3_)_2_, 695 μM KH_2_PO_4_, 60 μM K_2_HPO_4_, 20 μM FeSO_4_, 20 μM Na_2_EDTA, 9.25 μM H_3_BO_3_, 3.6 μM MnCl_2_, 74 nM (NH_4_)_6_Mo_7_O_24_, 3 μM ZnSO_4_, 785 nM CuSO_4_), pH is adjusted to 5.8. After 2 weeks of growth, plant roots were collected and stained with iodine. Roots were observed under phase contrast microscope (20X plan fluor, NA = 0.45, objective) and subsequently photographed.

### Protoplasts preparation and transformation

SS4 and PII1 cDNAs were cloned in a Gateway entry vector following manufacturer instructions (pENTR™ directional TOPO^®^ cloning cloning kit, Invitrogen). Using LR clonase (Gateway^®^ LR clonase™ II enzyme mix, Invitrogen), cDNA were transferred in the destination vector pUBC-GFP-Dest allowing expression of the protein fused to GFP (Grefen *et al*., 2010).

Arabidopsis protoplasts were isolated from 2-week-old plants grown on Murashige and Skoog agar (1.2 %) medium. Leaves were cut in 15 ml of 500 mM mannitol. After mannitol removal, preparations were incubated without shaking overnight in darkness at room temperature in enzyme solution (400 mM mannitol, 5 mM MES, 1 M CaCl_2_, 1% (w/v) cellulase Onozuka R10, 0.25 % (w/v) Macerozyme R10 at pH 5.6). Protoplasts were filtered through two layers of Miracloth (Calbiochem, EMD Biosciences, La Jolla, CA), centrifuged during 5 min at 50 *g* in swing out rotor at room temperature. Protoplasts were resuspended in 5 ml of W5 solution (154 mM NaCl, 125 mM CaCl_2_, 5mM KCl, 5 mM glucose, 1.5 mM MES, pH 5.6). 2.5 ml protoplasts were deposited on 6 ml 21 % sucrose solution and centrifuged 10 min at 50 g. Intact protoplasts, that accumulate on the sucrose surface were resuspended in 0.3 ml of MaMg solution (0.4 M mannitol, 15 mM MgCl_2_, 5 mM MES pH 5.6). Transformation was performed using 50 μg of plasmid DNA and 50 μg salmon sperm DNA used as sheared carrier DNA. 325 μl of transfection buffer was immediately added (40 % (w/v) PEG_4000_, 0.4 M mannitol, 0.1M Ca(NO_3_)_2_ pH7-8). Protoplasts were incubated for 30 min in darkness, washed in 10 ml of W5 solution, centrifuged 5 min at 50 *g* at room temperature and resuspended in 2 ml of W5 solution. Protoplasts were observed, 48-72 h after transformation, under a video microscope (Leica AF6000LX) with a Plan Apo 100x Oil (NA = 1.4) objective. We have observed protein expression with λ_ex_ = 484-500nm and λ_em_ = 514-554 nm (green channel), and protoplasts autofluorescence with λ_ex_ = 564-586nm and λ_em_ = 602-662 nm (red channel).

### Determination of starch granule number per chloroplast

One leaf of 2-weeks-old plants grown under 16 h : 8 h, light : dark photoperiod in a growth chamber was harvested at the end of the light phase and placed under vacuum in 1 ml fixating solution (4 % (w/v) paraformaldehyde, 4 % (w/v) sucrose, 1x PBS at pH 7.3). The leaf was then deposited between microscope slide and coverslip. Samples were observed under A1 Nikon confocal microscope (Nikon Instruments Europe B.V.) with a Plan Apo 60x Oil (NA = 1.4) objective. Auto fluorescence was acquired with λ_ex_ = 488 nm and λ_em_ = 500-550 nm (green channel), and with λ_ex_ = 561 nm and λ_em_ = 570-620 nm (purple channel).

### Polysaccharide extraction and purification

After 3 weeks of culture in a growth chamber, Arabidopsis leaves were harvested at the end of the day or at the end of the night. Samples were immediately frozen in liquid nitrogen, and stored at −80 °C until use. Depending on the subsequent analysis, two different extraction methods were performed.

For polysaccharide quantification we used a perchloric acid method (adapted from (Delatte *et al*., 2005)): About 0.3 g of leaves was homogenized with a Polytron blender in 5 mL of 1 M perchloric acid. The crude lysate was centrifuged at 4,500 *g* for 10 min at 4 °C to separate the pellet, which contains starch, and the supernatant containing the water soluble polysaccharides (WSP). The pellet was rinsed three times with sterile deionized water and resuspended in 1 ml H_2_O before quantification.

To determine starch granule size, polysaccharides chain length distribution profile and to perform scanning electron microscopy, starch was extracted as follow: Approximately 5 *g* of fresh material were homogenized using a polytron blender in 30 mL of the following buffer: 100 mM MOPS, pH 7.2; 5 mM EDTA; 10% (v/v) ethylene glycol. The homogenate was filtered through two layers of Miracloth (Calbiochem, EMD Biosciences, La Jolla, CA) and centrifuged for 15 min at 4 °C and 4,000 *g*. The pellet was resuspended in 2 ml Percoll 90% (v/v) and centrifuged for 40 min at 4 °C and 10,000 *g*. The starch pellet was washed with sterile distilled water (10 min at 4 °C and 10,000 g) and one time with 80% ethanol. Starch was finally stored at 4 °C in 20% ethanol.

### Granule size distribution and scanning electron microscopy

Purified starch granules were dispersed in 20 ml of IsoFlow Sheath (Beckman Coulter) and analyzed in a multisizer 4 Coulter-counter (Beckman) with a 20 μm aperture tube. The Multisizer software was set to determine the size of 30 000 particles ranging from 1 to 6 μm. 300 bins are logarithmically spaced between 1 and 6 μm (X-axis) and the size frequency distribution was expressed as relative percentage of total amount (Y-axis).

For scanning electron microscopy observation, drops of dilute aqueous suspensions of purified starch granules were deposited on a piece of glow-discharged copper tape and allowed to dry. The specimens were coated with Au/Pd and secondary electron images were recorded with an FEI Quanta 250 scanning electron microscope equipped with a field emission gun and operating at 2 kV.

### Starch content and ultrastructure

Leaves of 3-weeks-old plants were collected at the end of the day or at the end of the night and immediately frozen. Starch was extracted from 0.3 to 0.5g of leaves by the perchloric acid method described above. Starch content was determined by a spectrophotometric method following manufacturer’s instructions (Enzytec™ R-Biopharm). For each genotype, three independent cultures were performed. For each culture, three different samples were collected (each sample contains leaves from 3 plants).

Therefore, for each genotype, a mean and a standard error was calculated from nine different values (eight values for Col-O). A two-tailed *t*-test was applied to compare mutant lines to their respective wild-type.

The polysaccharide chain length distribution profile was determined on purified starch granules as described in (Boyer *et al*., 2016). 1 mg of purified starch was debranched with a mix of 4 U isoamylase (*Pseudomonas sp*, megazyme) and 2 U pullulanase (*Klebsiella planticora*, megazyme) in sodium acetate buffer (55 mM, pH 3.5) during 12 h at 42 °C. After desalting (Grace™ Alltech™ Extract-Clean™ Carbograph columns) and lyophilization, samples were suspended in 300 μl of deionized water. CLD was determined by high-performance anion exchange chromatography with pulsed amperometric detection analysis (Dionex® ICS-300 – PA200 CarboPac column 250×3 mm, Thermo Fisher, Sunnyvale, CA, USA) as fully described in (Roussel *et al*., 2013).

## Results

### *Selection of* pii1 *lines*

SS4 is a major protein involved in starch initiation. To identify other members of the starch-priming complex, proteins that potentially interact with SS4 were identified by a yeast-2-hybrid screen. An ULTImate Y2H™ was carried out by Hybrigenics-services (Paris, France) using SS4 as bait (amino acids 43 to 1040) against a library prepared from one-week-old seedlings. Among 125 millions interactions tested, 369, corresponding to 80 different proteins, were positive. During this screen, SS4 was identified in the prey proteins, confirming that it is able to interact with itself and validating the approach. Using ChloroP algorithm prediction (Emanuelsson *et al*., 1999) on SS4-interacting candidates, we have selected proteins with predicted chloroplast targeting peptides. Among six SS4-interacting selected candidates having a predicted chloroplast targeting peptide (Table S1), we have identified the PII1 protein (*At4g32190*). For each interaction identified during the screen a predicted biological score (PBS) was given. This score indicates the reliability of the identified interaction and ranges from A (very high confidence of the interaction) to F (experimentally proven artifacts). The protein encoded by the gene *At4g32190* displayed a PBS of “C” corresponding to a “good confidence of interaction”. Indeed, SS4 used as bait was found to interact with 5 yeast clones expressing different fragments of the pray protein. An *in vivo* interaction between SS4 and PII1 was also observed by bimolecular fluorescence complementation (BiFC) experiments. Tobacco plants were transformed to allow the transient co-expression of SS4 and PII1. Each protein was fused to a moiety of YFP. The fluorescence corresponding to YFP reconstitution reveals the proximity of SS4 and PII1 in tobacco chloroplasts (Supporting information Fig. S1).

To determine the biological significance of the potential interaction between this protein and SS4, we have engaged a phenotypic analysis of lines impaired in the corresponding gene. Two Arabidopsis lines with T-DNA insertion within *At4g32190* gene were obtained from NASC resource center. Lines N679037 and 137A02 were in Col-0 and Ws genetic background respectively. After selection of homozygous lines, we have evaluated *At4g32190* mRNA integrity by RT-PCR (Fig. 1). In both cases the RNA integrity is compromised by the T-DNA insertion. In line N679037, the insertion is located within 5’ UTR (three nucleotides before the ATG codon), while in line 137A02, T-DNA is inserted in the second exon. The two mutant alleles were named *pii1-1* (Col-0 background) and *pii1-2* (Ws background), respectively.

**Fig. 1:**
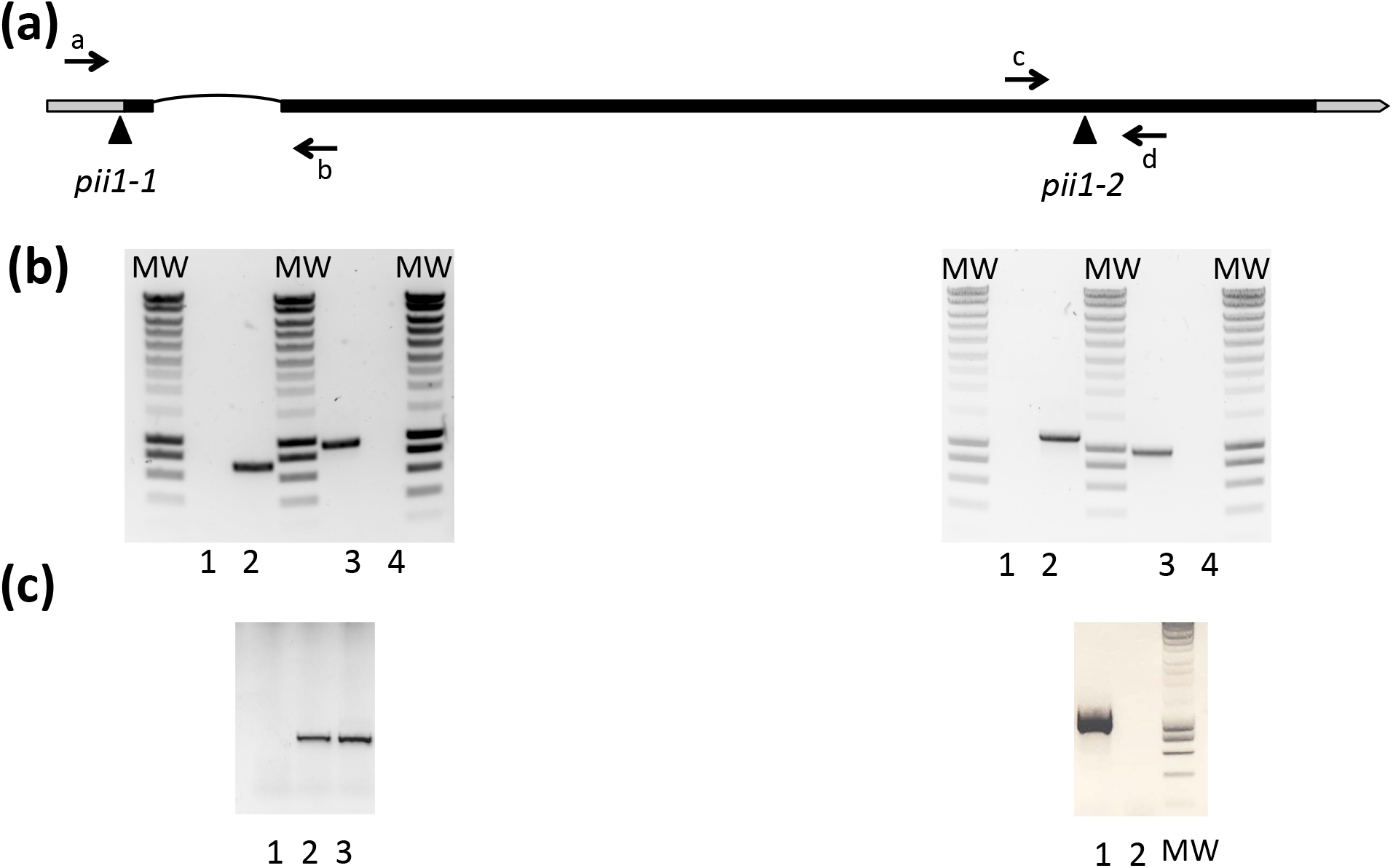
Selection of *pii1* mutants. (**a**) Structure of the *At4g32190* locus encoding PII1. UTR are indicated by grey boxes while introns and exons are depicted as black lines and black boxes respectively. T-DNA insertion corresponding to *pii1-1* and *pii1-2* alleles are indicated by triangles. Primer position used for selection are indicated by arrows (not in scale) (**b**) Left panel represents the selection of the *pii1-1* homozygote mutant. gDNA from one mutant plant (lanes 1 and 3) and one wild-type control (lanes 2 and 4) was used. Amplification products of wild type allele (using primer a and b) are in lanes 1 and 2. Amplification products of mutant allele (using T-DNA primer and primer b) are in lanes 3 and 4. Right panel represents the selection of the *pii1-2* homozygote mutant. gDNA from one mutant plant (lanes 1 and 3) and one wild-type control (lanes 2 and 4) was used. Amplification products of wild type allele (using primer c and d) are in lanes 1 and 2. Amplification products of mutant allele (using T-DNA primer and primer c) are in lanes 3 and 4. (**c**) Left panel: RT-PCR amplification products obtained using primers a and b on total RNA extracted from *pii1-1* (lane 1), *ss4-1* (lane 2) and Col-0 (lane 3). Right panel: RT-PCR amplification products obtained using primers c and d on total RNA extracted from Ws (lane 1), *pii1-2* (lane 2). Molecular weight (MW): SmartLadder, Eurogentec.

Even if PII1 is annotated as a « myosin heavy chain-related protein » in databases, no function has been reported for this protein. Analysis performed using MARCOIL server (Delorenzi & Speed, 2002; Zimmermann *et al*., 2017) indicates that PII1 is composed of several coiled-coil domains. Considering only the amino acids that have a probability above 50% to be involved in a coiled–coil motif, four distinct domains are identified (regions 106-331; 340-426; 448-529 and 629-723). Altogether these motifs represent more than 60% of the protein sequence. The coiled-coil motifs are known to favor protein-protein interaction (Adamson *et al*., 1993) and interestingly, it has already been reported that SS4 also contains such motifs (Leterrier *et al*., 2008).

Phytozome server allowed the identification of Arabidopsis PII1 homologs in other dicots, such as *Solanum tuberosum* (XP_006344374) and in monocots (ex: *Brachypodium distachyon* XP_010233922; *Oryza sativa:* XP_015627751; *Zea mays:* XP_008679905). Homologs can also be found in lycopodiophytes (*Selaginella moellendorffii*: XP_002983651), but no protein similar to PII1 was identified in bryophytes (*Physcomitrella patens*) or green algae such as *Chlamydomonas reinhardtii* or *Ostreococcus lucimarinus*. Interestingly no clear homologs of SS4 can be found in these green algae indicating that a co-evolution of PII1 and SS4 may have occurred.

### Intracellular localization of the PII1 protein

*At4g32190* encodes a protein of 783 amino acids. Use of ChloroP software (Emanuelsson *et al*., 1999) predicts a chloroplast targeting peptide of 27 amino acids in length. Even if the size of this predicted targeting peptide is relatively small, it is not incompatible with the predicted subcellular localization (Bionda *et al*., 2010). Nevertheless PII1 localization was experimentally determined by protoplast transient transformation. Protoplasts were prepared from Arabidopsis wild-type or *pii1* plants and transformed with a construct allowing the transient expression of PII1 fused in its C-terminus to the green fluorescent protein (GFP). Images obtained using a fluorescence microscope confirm the chloroplastic localization of the protein. Moreover fluorescence appears as dots (Fig. 2), a pattern that is similar to that already reported for SS4 (Raynaud *et al*., 2016). Protoplasts of the *ss4* mutant were also transformed with the PII1-GFP fused chimeric construct. Again, the PII1-GFP protein was located within chloroplasts and the same dot-like distribution pattern of the protein was maintained indicating that SS4 is not required for proper PII1 localization inside the chloroplast.

**Fig. 2:**
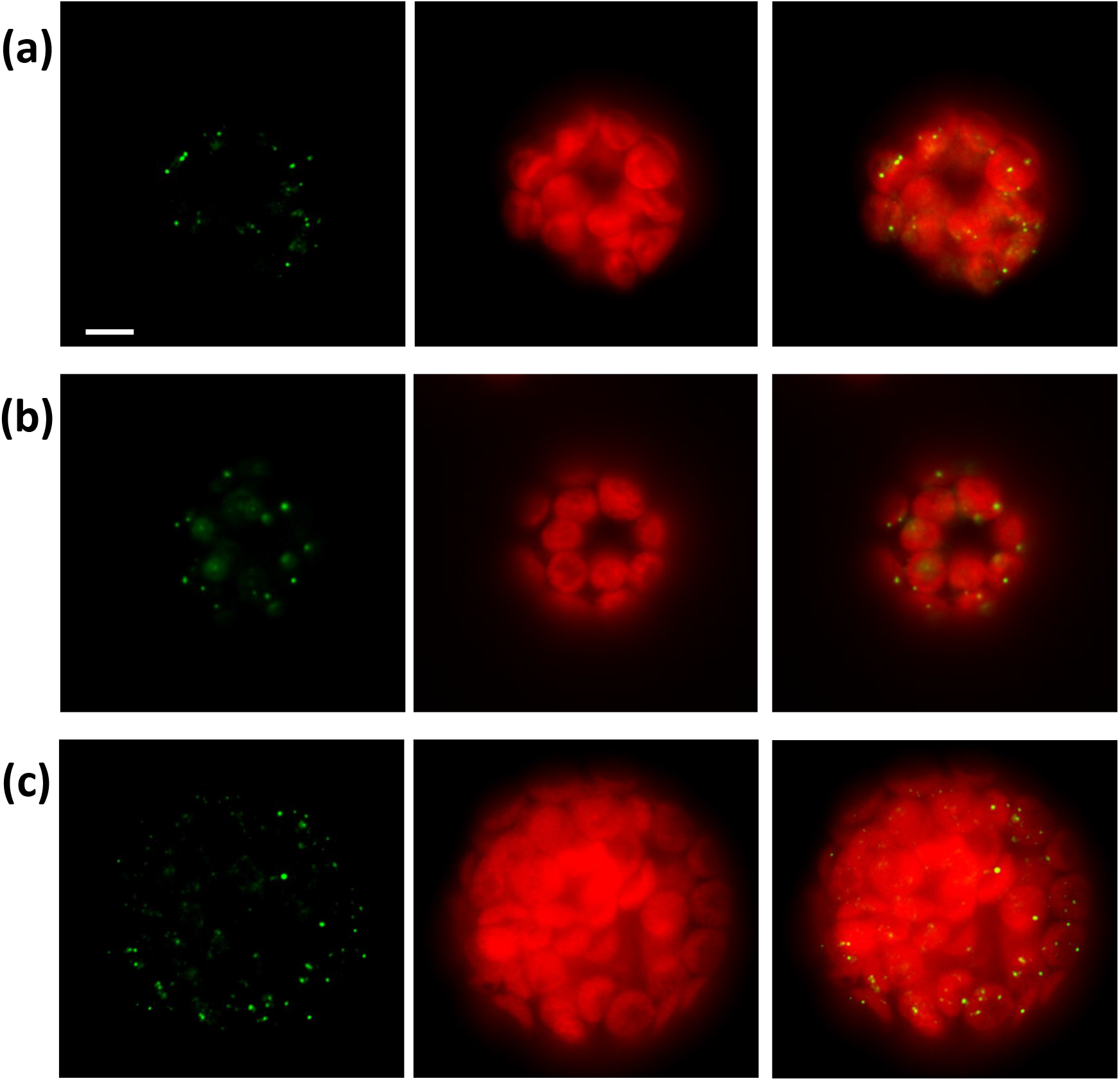
Subcellular localization of PII1 in Arabidopsis protoplasts. Protoplasts were prepared from Col-0 (**a**), *pii1-1* (**b**) or *ss4-1 pii1-1* (**c**). Image acquisition was performed using a video microscope. In each row, the first image corresponds to the GFP signal of the PII1-GFP fusion protein. Chlorophyll fluorescence is displayed in the central panels. The merged images are displayed on the right. Scale bar = 10 μm

### Starch granule number and morphology are altered in pii1 lines

The starch granule number per chloroplast was determined from light microscopy observations of Arabidopsis leaf cells. This technique allows the visualization of starch granules without the need to produce sections of the leaf tissue. While a typical number of 5 to 7 starch granules were observed in the chloroplasts of wild-type cells, most plastids of *pii1* leaf cells contained only one large starch granule (Fig. 3). This observation was made whatever the genetic background of *pii1* lines (i.e., Ws or Col-0) indicating a reduction of starch-granule priming efficiency in these mutants.

**Fig. 3:**
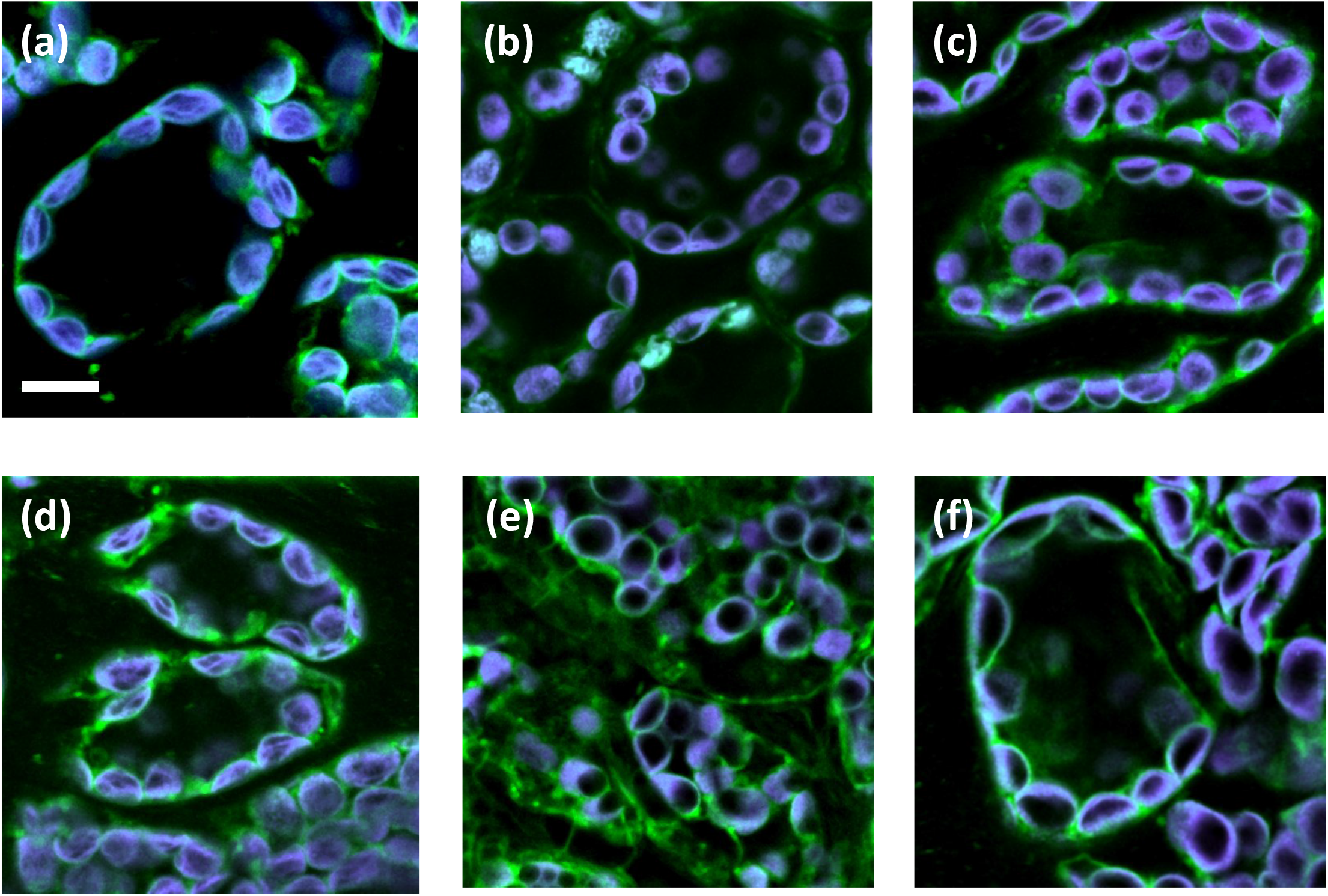
Starch granule number per chloroplast. Isolated leaf cells were prepared from Col-0 (**a**); *ss4-1* (**b**); *pii1-1* (**c**); Ws (**d**); *ss4-2* (**e**) and *pii1-2* (**f**). Leaves samples were collected at the end of the day and tissues were fixed in paraformaldehyde before disruption in EDTA. Pictures were collected using a confocal microscope. Chlorophyll fluorescence in purple delimits the chloroplast volume, while starch granules are black. Scale bar = 10 μm

The granule size was determined from starch extracted at the end of the day from the rosette leaves of 3-week-old plants. The use of a Coulter counter allows the determination of 30,000 particles size within few seconds. In our growth conditions (16 h : 8 h, light : dark photoperiod), wild-type (WT) starch displays an unimodal distribution of granules size with a maximum at 1.5 and 1.6 μm for Col-0 and Ws ecotypes, respectively. As already reported, starch granules of *ss4* lines are larger with a maximum of the Gaussian distribution between 3.4 and 3.5 μm (Fig. 4). Starch granules extracted from *pii1* lines are also larger compared to the WT counterpart. Distributions of granule size remain unimodal in both mutant lines with a peak of the Gaussian distribution at 2.5 and 3.3 μm in Col-0 and Ws background, respectively (Fig. 4).

**Fig. 4:**
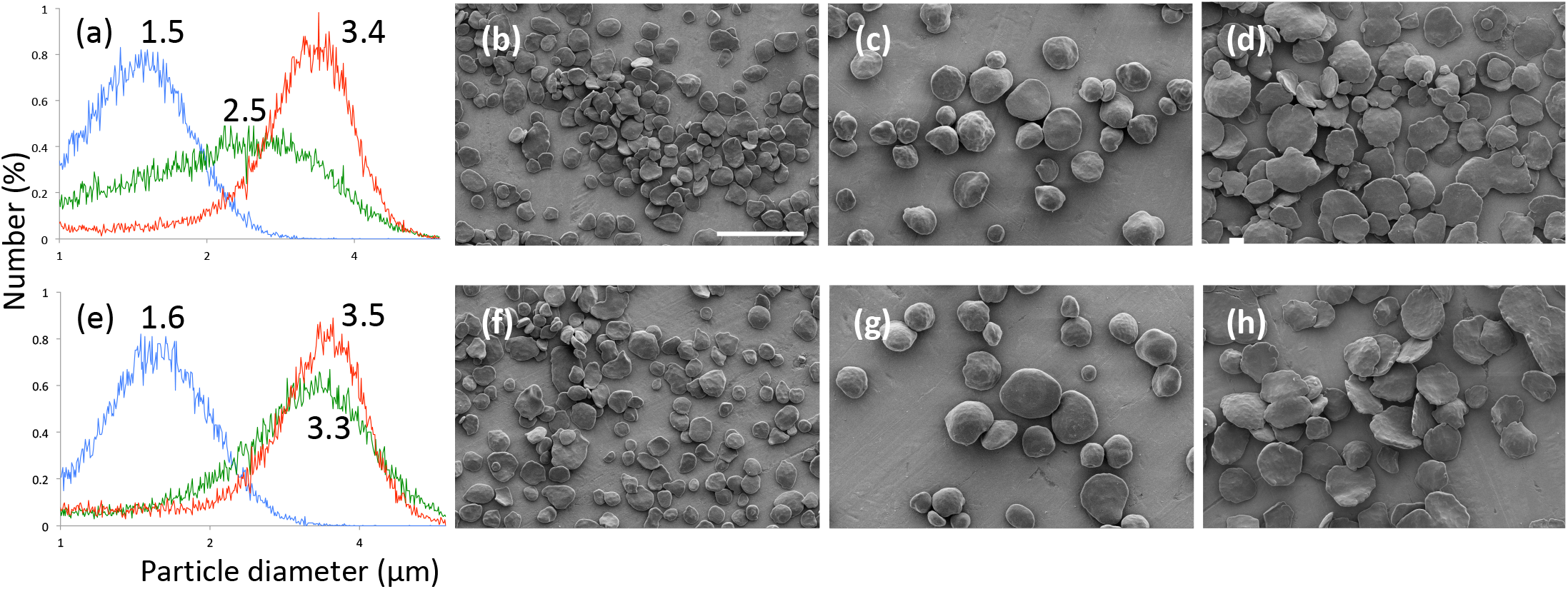
Starch granule size and morphology. Starch granules were extracted and purified from leaves of 3-week-old plants harvested at the end of the day. The plants were grown in 16 h : 8 h, light : dark photoperiod. Panels (**a**) − (**d**): plants of the Col-0 genetic background. Panels (**e**) − (**h**) plants of the Ws genetic background. The starch granule size distribution was determined by analyzing 30,000 particles extracted for each genotype. The results are expressed in relative percentage (*y*-axis) of particles of a diameter ranging from 1 to 6 μm (x-axis, logarithmic scale). In (**a**) and (**e**), starch from wild type, *ss4* and *pii1* lines are in blue, red and green, respectively. The value at the peak of the Gaussian distribution is indicated in μm. (**b**) to (**d**) and (**f**) to (**h**): starch granules were observed using scanning electron microscopy. (**b**) and (**f**): wild-type (Col-O and Ws, respectively); (**c**) and (**g**): *ss4-1* and *ss4-2*; (**d**) and (**h**): *pii1-1* and *pii1-2*. Scale bar = 10 μm, all images are at the same scale.

The morphology of purified starch granules was also examined using scanning electron microscopy. The granules extracted from wild-type Arabidopsis leaves were lenticular in shape with a smooth surface. Several mutations affecting Arabidopsis starch-metabolizing enzymes were reported to affect starch granule morphology (Szydlowski *et al*., 2011; Malinova & Fettke, 2017). This is also the case of the *ss4* mutants that accumulate large and spherical granules with a smooth surface. While similar in size to granules extracted from *ss4* mutants, those purified from *pii1* remain lenticular but display an indented surface (Fig. 4).

### Impact of PII1 deficiency on plant growth and root development

*ss4* mutants display growth retardation and pale green leaves phenotype. These characteristics were proposed to be caused by the reduction of initiation events leading to accumulation of unused ADP-glucose especially in starch free chloroplasts (Ragel *et al*., 2013). Since *pii1* mutants lines also display a reduction of initiation events, growth kinetics of these lines were analyzed and compared to wild-type and *ss4* lines. Plants were grown in a green house under 16 h light and 8 h dark cycling conditions during 3–weeks. Growth rate of *pii1* lines was similar to wild type while growth retardation of *ss4* lines was confirmed whatever the genetic background (Fig. 5). Pale green leaves phenotype was recorded only in SS4 deficient lines.

**Fig. 5:**
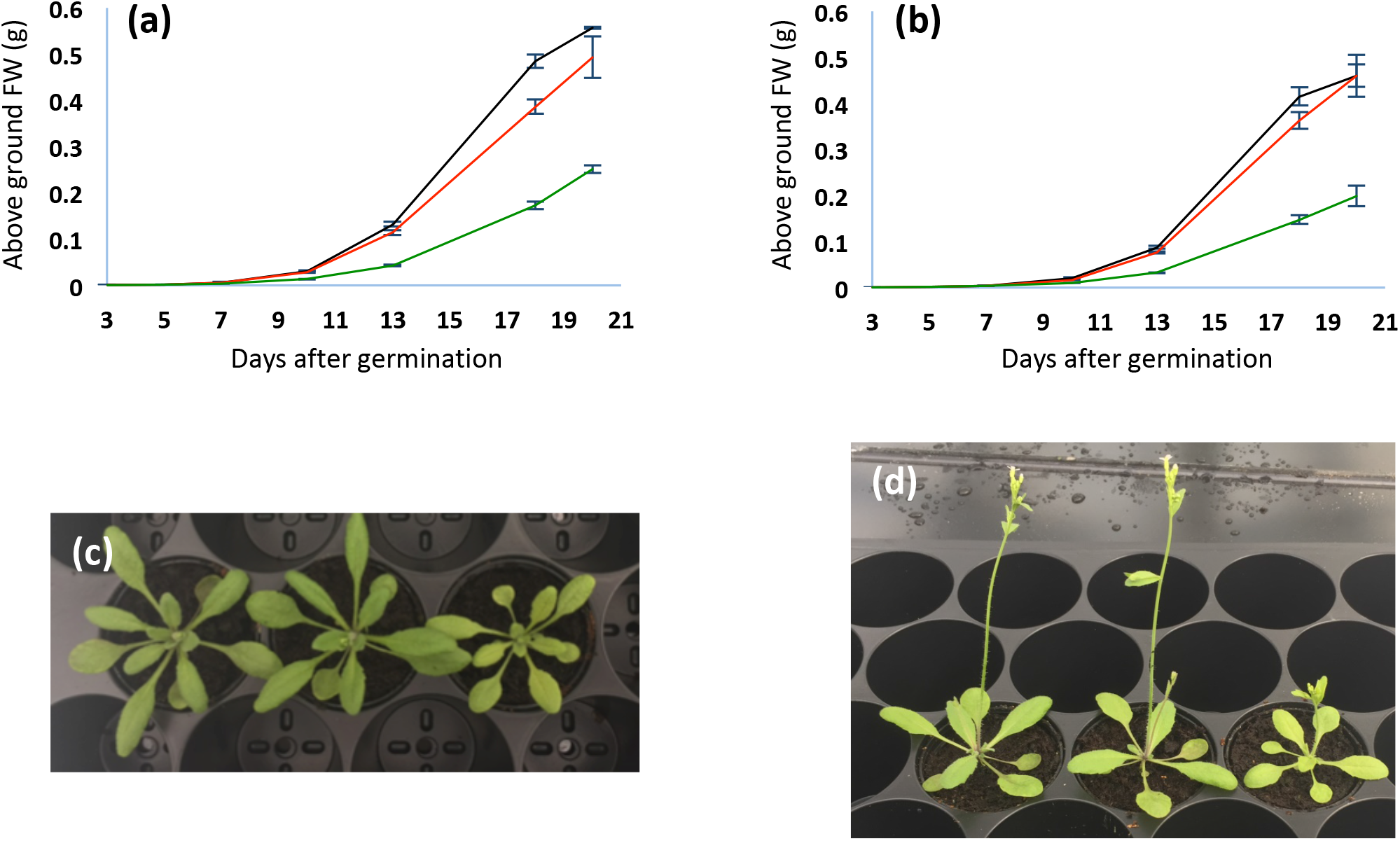
*piil* growth phenotype compared to *ss4* and wild type plants. Plants were grown in a greenhouse under 16 h : 8 h, light : dark photoperiod. Panels (**a**) and (**b**): the weight of above ground organs (g/plant) is plotted against the number of days after germination. Black lines correspond to wild type plants (Col-0 and Ws in panels (**a**) and (**b**) respectively), red lines correspond to *pii1-1* and *pii1-2* in panels (**a**) and (**b**) respectively and green lines correspond to *ss4-1* and *ss4-2* in panel (a) and (b) respectively. Each value corresponds to the mean of three samplings (each sample being composed of several plants). Thin vertical bars represent the standard error. Panels (**c**) and (**d**) are pictures of 3-week-old plants grown in the same conditions as above. Panel (**c**): from left to right: Col-0, *pii1-1, ss4-1*. Panel (**d**): from left to right: Ws, *pii1-2, ss4-2*.

Starch accumulation in roots and roots development were also evaluated in *pii1* mutants (Fig. 6). Plants were cultivated in a growth chamber using hydroponic systems and starch accumulation in the columella of primary and lateral roots was visualized under microscope after iodine staining. The alteration of starch synthesis at the apex of *ss4* roots was previously described and this alteration was associated with a modification of root architecture and response to gravity (Crumpton-Taylor *et al*., 2013). After iodine staining of 2-weeks-old *pii1* seedlings, the apex of both primary and lateral roots appear dark blue indicating that *pii1* mutants accumulate starch granules in the columella. The intensity of the coloration and number of stained granules in *pii1* appear to be similar to wild type. Moreover, the architecture of root development was visualized on agar plates that were placed vertically after seed germination. Again, the *pii1* line behaves like wild-type plants and the roots of 2-week-old seedlings respond correctly to gravity and no aberrant development was observed (Fig. 6).

**Fig. 6:**
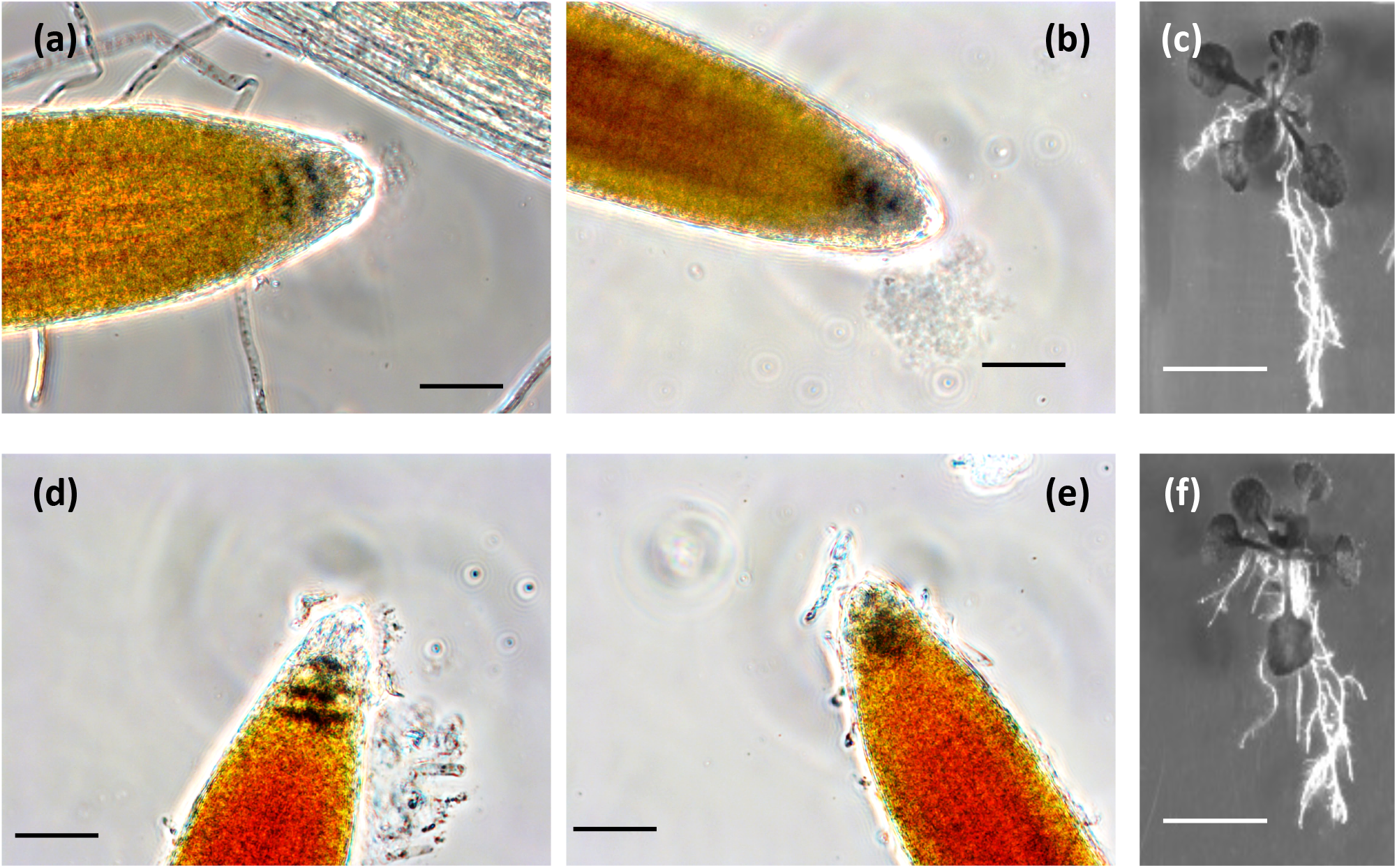
Starch accumulation in root and root development. Plants were grown during 2 weeks under 16 h : 8 h, light : dark photoperiod. Panels (**a**), (**b**) and (**c**): Ws. Panels (**d**), (**e**) and (**f**): *pii1-2*. Panels (**a**), (**b**), (**d**) and (**e**): roots of 2-week-old plants cultured with hydroponic systems were soaked in lugol, rinsed with water and observed under microscope. (**a**) and (**d**): primary root. (**b**) and (**e**): lateral root. Panels (**c**) and (**f**): seeds were sown on agar plates that were maintained vertically after seed germination. Pictures were taken after 2 weeks of growth. Black bar = 50 μm; white bar = 1 cm

### Impact of PII1 deficiency on starch amount and ultrastructure

The mutation at the *PII1* locus led to the increase of starch granule size associated to a reduction of granule number. To determine if these phenotypes are associated to a modification of total starch amount, we also assayed the starch content in these lines and compared data to wild-type and *ss4* lines. Leaves of 3-week-old plants, grown in a growth chamber under 16 h : 8 h, light : dark photoperiod, were harvested at the end of the illuminated or dark periods and the starch content was enzymatically assayed. It was similar to the amount already reported for wild-type and SS4-deficient lines (Roldán *et al*., 2007). The loss of SS4 led to a slight reduction of starch content at the end of the day but a higher amount at the end of the night (Fig. 7). This effect is exacerbated in Ws background. In *pii1* mutants, no reduction of the starch content was recorded at the end of the day. Moreover, in Ws genetic background, the residual starch accumulated at the end of the night was significantly higher in *pii1* or *ss4* mutants compared to the corresponding wild-type line (Fig. 7). No significant accumulation of water-soluble polysaccharides could be detected in the different analyses.

**Fig. 7:**
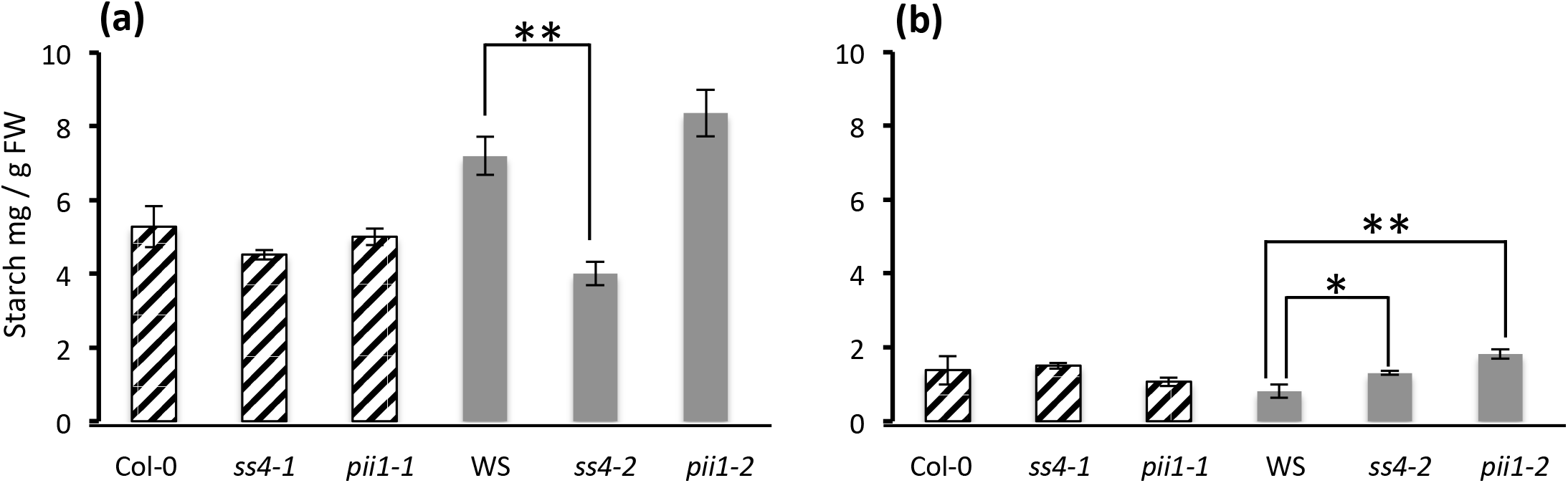
Starch content in leaves. Plants were grown in a culture room under 16 h : 8 h, light : dark photoperiod. Three weeks after germination leaves were harvested either at the end of the day (**a**) or at the end of the dark phase (**b**). Three independent cultures were performed and for each culture three independent extractions were realized. Values correspond to the mean of nine assays (except for Col-0 end of dark phase corresponding to 8 assays). Thin vertical bars represent the standard error. Values obtained for mutant lines were compared to their respective wild-type by a two-tailed *t*-test. Asterisk represents statistically significant difference at p < 0.05 (*) or p < 0.001 (**)

Starch fractionation was carried out by size exclusion chromatography on Sepharose CL-2b matrix. No modification of the amylose / amylopectin ratio was recorded in *pii1-1 or pii1-2* lines when compared to their respective wild types (Supporting information Fig. S2). The starch ultrastructure was also analyzed by establishing the chain length distribution (CLD) profile of the glucans. The purified starch was enzymatically debranched and linear glucans were then separated using high-performance anion exchange chromatography and detected by pulsed amperometric detection (HPAEC-PAD). The CLD profile determined for starch extracted from PII1 deficient lines was identical to that of the corresponding wild types (Fig. 8).

**Fig. 8:**
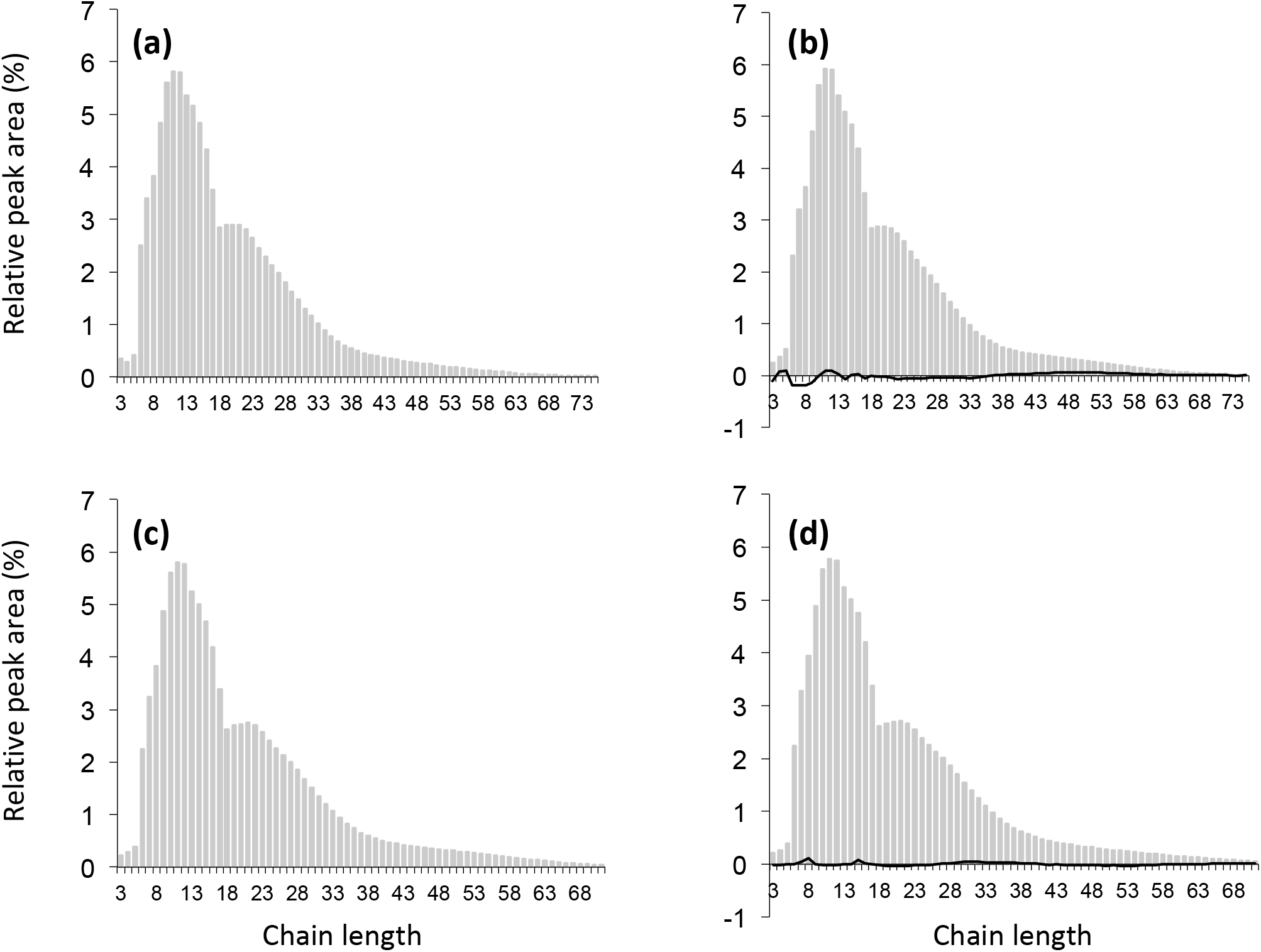
Chain length distribution of purified starches. Purified starches were enzymatically debranched. Linear glucans were then separated and detected using high performance anion exchange chromatography with pulsed amperometric detection (HPAEC-PAD). Grey bars represent the proportion of each DP expressed as a percentage of the total amount presented in the figure. The black line (panels (**b**) and (**d**)) corresponds to the differential plot between the mutant and wild type profiles. (**a**): Col-0. (**b**): *pii1-1*, (**c**): WS. (**d**): *pii1-2*. Each profile is the mean of two analysis carried out with starch extracted from two independent cultures.

### PII1 deficiency does not alter other starch metabolizing enzymes

The activity level of several enzymes was estimated by zymogram analysis. All tested activities (starch synthases SS1 and SS3, branching enzymes, debranching enzymes, phosphorylases) remained unaffected in the *pii1* mutant (Supporting information Fig. S3).

Since the reduction of granules number without alteration of starch structure observed in *pii1* line was similar to that observed in *ss4, ptst2* or *ptst3* lines, we verified that these genes were still correctly expressed in the absence of PII1. Total RNAs of *pii1* lines were extracted and purified from leaves harvested in the middle of the day. RT-PCR performed to amplify RNA signal of SS4, PTST2 and PTST3 gave signals in *pii1* lines indicating that these genes are expressed (Supporting information Fig. S4). In the same manner, the RNAs were extracted and purified from *ss4* lines and RT-PCR revealed that PII1 is expressed in this mutant (Fig. 1).

Moreover, SS4 was reported to be located in specific regions of the chloroplast. SS4 is preferentially distributed near the thylakoid membranes (Gámez-Arjona *et al*., 2014). The N-terminal moiety of SS4, containing several coiled-coil domains known to be involved in protein-protein interactions, is responsible of the proper location of SS4 (Lu *et al*., 2018). To determine if PII1 has an impact on SS4 subchloroplastic location, we expressed the SS4-GFP fused protein in the *pii1* mutant. SS4 distribution was not modified in the presence or absence of PII1 (Fig. 9).

**Fig. 9:**
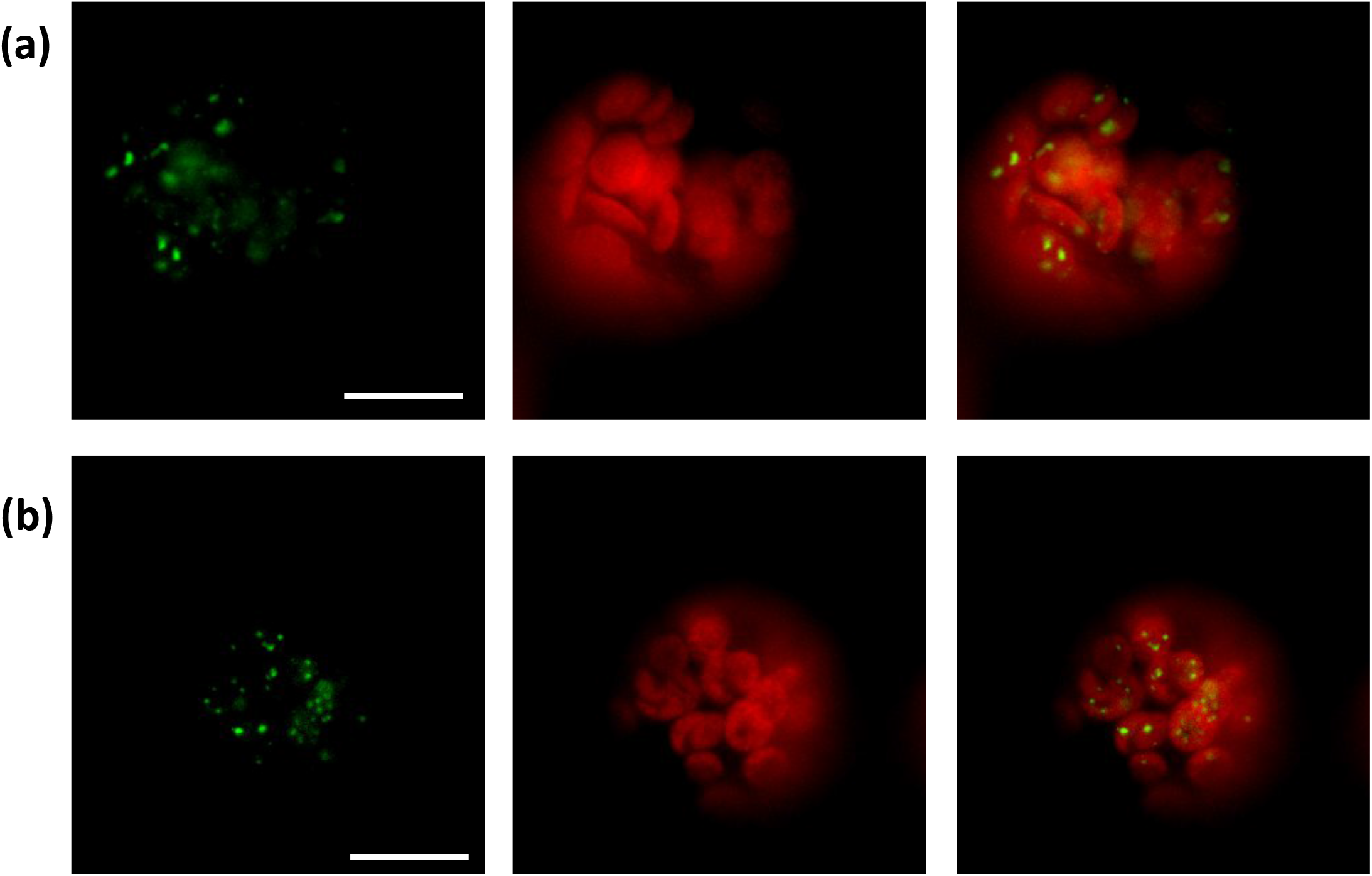
Localization of SS4 in Arabidopsis protoplasts. Protoplasts were prepared from *ss4-2* (**a**), *ss4-2 pii1-2* (**b**). Image acquisition was performed using a video microscope. In each row, from left to right: SS4-GFP fluorescence; Chlorophyll fluorescence; Merged images. All images are at the same scale. Bar = 10 μm

## Discussion

### PII1 and SS4 interact in planta

In this study we describe a new protein involved in starch priming. PII1 was identified by yeast-2-hybrid screen as physically interacting with SS4. The number of positive clones with different PII1 fragments gave a good confidence of the interaction between the two proteins. This result was reinforced by other observations such as subcellular protein localization or BiFC experiment.

*In silico* analysis of PII1 coding sequence predicts the presence of a small transit peptide allowing the targeting of the protein to the chloroplast (Table S1). The chloroplastic localization of PII1 was confirmed by expression of the recombinant PII1-GFP fluorescent chimeric protein. Moreover this approach also revealed a dotted distribution of the protein in the plastid (Fig.2) similarly to what was previously described for SS4 (Raynaud *et al*., 2016). Although the formation of aggregates of PII1 could not be totally excluded, it is unlikely that such dotted distribution of PII1 arises from the over-expression of the protein, since we have made use of the moderate expression “ubiquitin promoter-10” for the expression of the transgene (Grefen *et al*., 2010). In addition, the transformations were carried out with protoplasts prepared from cells of the *pii1* mutant lacking the endogenous protein limiting the over accumulation of the protein. Finally, the spatial proximity of PII1 and SS4 has also been confirmed *in planta* by a BiFC approach (Supporting information Fig. S1).

As a whole, these results (yeast-2-hybrid, distribution within the chloroplast and BiFC) provide good evidence for the direct physical interaction between SS4 and PII1 in the chloroplast.

### PII1 is required in determination of starch granule number in Arabidopsis chloroplast

In this work, we have phenotypically characterized two independent mutant alleles of the PII1 gene: the *pii1-1* in the Col-0 background and *pii1-2* in the Ws background. T-DNA insertion lies in the 5’UTR, only 4 nucleotides upstream of the START codon in *pii1-1*, while it is localized in exon 2 in *pii1-2*. Although these two lines were confirmed mutant by RT-PCR (Figure 1) we cannot exclude that a small amount of active PII1 is still produced at a low level in the *pii1-1* mutant while it is completely abolished in *pii1-2*.

Nevertheless, the phenotypes of the two mutants are very similar. Both lines have less (typically one) but bigger starch granules per chloroplast. Although the effect is more pronounced in *pii1-2* (*pii1-1* is probably not a null allele), starch granules of PII1 deficient lines are 2 to 3 times larger than wild type granules, and of a size similar to those of the *ss4* mutant. However, while *ss4* starch is spherical in shape, those extracted from *pii1* mutants remain lenticular, a form comparable to that of wild-type starch.

Another difference observed between the *pii1* and the *ss4* mutants is related to plant growth. Indeed, while the *ss4* mutants show significant growth retardation, the *pii1* behave like their corresponding wild types. The stunting growth of *ss4* seems, at least partly, related to the accumulation of higher ADP-glucose content in the leaves (Ragel *et al*., 2013) probably because a significant fraction of chloroplasts are starch-free in these mutants (Roldán *et al*., 2007; Lu *et al*., 2018). Although the absence of PII1 leads to a reduction in the number of starch granules per chloroplast, none or very few organelles are completely free of starch (a much smaller fraction than ss4). In addition, the *ss4* mutants have a root growth defect related to the perturbation of starch accumulation in the columella (Crumpton-Taylor *et al*., 2013). Starch accumulation in roots and root development of *pii1* lines are similar to wild type plants. The combination of these two factors, absence of starch-free chloroplast and normal root development, may explain the correct growth of plants lacking PII1.

### PII1 is proposed to be required for SS4 catalytic activity

SS4 is a major component involved in starch priming. This protein is composed of two distinct parts. The C-terminal part of the protein corresponds to the catalytic domain that is shared by all starch-synthases of the GT5 family. The N-terminal part of SS4, essentially composed of coiled-coil domains, is unique (Raynaud *et al*., 2016). Both parts have specific function in determination of starch granule size and morphology. Expression in *ss4* mutants of a truncated protein lacking the N-terminal part or a chimeric protein composed of N-terminal part of SS4 fused to the glycogen-synthase demonstrates that the catalytic domain (C-terminal) of SS4 determines the number of starch granule, while the N-terminal moiety control the shape of the granule (Lu *et al*., 2018). In the present work, PII1 was identified as an SS4-interacting partner. When PII1 was knocked down, the major phenotype was the reduction of starch granule number per chloroplast without alteration of granule shape. This phenotype can be interpreted as an inactivation of SS4 catalytic part (C-terminal) without alteration of the function of SS4 N-terminal moiety. We therefore propose that PII1 is a protein involved in starch priming that interacts with SS4 and is required for proper catalytic activity of the starch synthase controlling the number of initiation events generating new starch granules. PII1 could be needed to provide an adequate substrate to SS4 and / or to prevent the degradation of the substrate presented to SS4 to prime granule formation. A similar function was already proposed for PTST2 another SS4-interacting protein (Seung *et al*., 2017). Alternatively, PII1 could be requested to the correct folding of SS4 or its association with any other factor requested in the starch-priming machinery. Leaves are submitted to diurnal fluctuations of the starch content. The precise control of the starch content is of prime importance in leaves to prevent starvation at night (Scialdone *et al*., 2013). Such precise control could only be achieved if the priming of starch synthesis (number, size, and morphology of the granules) is itself highly controlled. Consequently, starch granule initiation is a complex mechanism involving enzymes and non-catalytic proteins. To date, several components of this machinery have been identified including SS4, SS3, PTST2, PTST3 and the new PII1 protein described in this work. Further investigations are needed to determine the precise function of each protein within the starch-priming complex and understanding how plants control the number and the size of the starch granules.

## Acknowledgments

The authors thank Christine Lancelon-Pin (CERMAV, Grenoble) for the SEM observations and the NanoBio-ICMG platform (FR 2607, Grenoble) for granting access to the Electron Microscopy facility.

We are indebted to the Research Federation FRABio (Univ. Lille, CNRS, FR 3688, FRABio, Biochimie Structurale et Fonctionnelle des Assemblages Biomoléculaires) for providing the scientific and technical environment conducive to achieving this work

We thank the Agence Nationale de la Recherche for funding (Project “CaSta-DivA” Programmes non thématiques 2011 section SVSE6).

## Author Contribution

CV performed most experiments. AC, DD, MF participated to experiments. AW performed chain length distribution analysis. JLP has supervized the electron microscopy observations, CS performed optical microscopy observations. FW and CDH designed experiments and wrote the manuscript.

## Supporting information

**Table S1**: Proteins identified during the yeast-2-hybrid screen, that are predicted to be targeted to the chloroplast.

**Table S2**: Information on T-DNA lines and primers used for selection and RT-PCR experiments.

**Fig. S1**: SS4 and PII1 interaction (BiFC).

**Fig. S2**: Starch fractionation.

**Fig. S3**: Zymograms of starch metabolizing enzymes.

**Fig. S4**: expression of *SS4, PTST2* and *PTST3*.

